# Protection of savings by reducing the salience of opposing errors

**DOI:** 10.1101/2024.05.03.592370

**Authors:** Mousa Javidialsaadi, Scott T. Albert, Badr Moufarrej S Al Mutairi, Jinsung Wang

**Affiliations:** Kinesiology programs, Zilber College of Public Health, University of Wisconsin–Milwaukee, Milwaukee, WI, 53201, USA; Neuroscience Center, University of North Carolina at Chapel Hill, Chapel Hill, NC 27514, USA

**Keywords:** Gradual adaptation, Visual rotation, Washout, Unlearning, Implicit learning, Readaptation

## Abstract

When humans encounter the same perturbation twice, they typically adapt faster the second time, a phenomenon called savings. Studies have examined savings following adaptation to a gradually introduced perturbation, with mixed results. These inconsistencies might be caused by differences in how behavior returns to its baseline state during the ‘washout’ phase in between learning periods. To test this, participants controlled a cursor that was subject to a visual rotation in its motion direction. The rotation was applied during two learning periods, separated by a washout period where the rotation was removed abruptly, gradually, or without error feedback. We found that the type of error experienced during washout affected savings: abrupt washout with large errors eliminated savings, whereas gradual or no-feedback washout preserved it. Model-based analyses indicated these effects were driven by changes in error sensitivity suggesting that salient, opposing errors experienced during washout downregulate the response to error, nullifying savings.

## Introduction

When an unexpected perturbation to our motor system occurs, the resulting errors in our actions and predictions^1–5^ update our motor commands, predictively countering future disturbances. For example, when we walk on slippery ice, a sudden fall results in an unwanted error that alters how we traverse the icy patch in the future. Even when the icy patch (i.e., the perturbation) is removed and we reestablish our normal walking patterns, a lingering memory is stored within the nervous system, whereby re-exposure to the perturbation results in more rapid re-learning. This learning hallmark, termed savings^6^, can occur on the same day, overnight^7^, or even a year after the initial perturbation^8^, and is ubiquitous across movements executed with our eyes (saccade adaptation^9^), legs (locomotion^10,11^), arms (visuomotor^7,12,13^ and force field adaptation^14–16)^, as well as classical conditioning paradigms^17^.

Our ability to flexibly increase our learning rate is known to rely on two parallel adaptive systems: an explicit process directed by our conscious strategies^18–20^, and a subconscious implicit process that operates without our awareness^2,4^. The implicit and explicit contributions to savings in motor tasks are typically studied using visuomotor rotations. In this task, subjects control some type of cursor with their hand or arm and move it to a target location. Following the baseline period of task familiarization, a visuomotor perturbation is introduced that rotates the cursor’s trajectory relative to the hand’s actual reaching direction. This is akin to altering the orientation of your laptop so that moving your cursor up results in an unexpected diagonal trajectory. The visual rotation thus induces an error, which individuals use to alter their behavior over time^21^. This altered behavior, or in this case, a preemptive change in the direction you move your hand to counteract the cursor’s rotation, is supported by intentional changes in the aiming direction you select (i.e., explicit strategy) along with an additional uncontrolled adjustment (*i.e.*, the implicit correction)^18^.

In visuomotor rotation tasks, improvements in strategy use are salient in cases where a large rotation is introduced abruptly. The abrupt introduction of a large rotation causes large errors in the anticipated cursor motion, which in turn encourages individuals to adopt an intentional re-aiming strategy^22–24^. If the rotation is reintroduced in the future, participants recall their past strategy, resulting in a greatly accelerated re-learning rate (termed “explicit savings”)^7,12,25^. This form of explicit savings can be suppressed either by limiting response time^22^ (which causes people to respond reflexively to the presentation of a target, thereby preventing them from using a re-aiming strategy) or by reducing the error magnitude during initial learning^26^ (which reduces awareness of the rotation). The latter is typically done by gradually introducing the rotation. When the rotation size increases gradually, our slow-adapting, subconscious (i.e., implicit) learning system is better able to mitigate the cursor errors, resulting in smaller discrepancies between intended and actual cursor movements. This can help prime the implicit system to exhibit savings that are detectable during subsequent encounters with the rotation^22,26^.

Intriguingly, however, studies that employed gradual rotations have observed mixed effects on the ability of *gradual* learning to facilitate more rapid *abrupt* re-learning. Absence of savings following gradual learning^27^ is consistent with a standard memory of errors model^27,28^, in which gradual learning cannot improve abrupt learning due to a mismatch in error size: errors are small in the gradual case, and thus gradual learning fails to generalize to abrupt learning in which large errors are experienced. Thus, to the brain, an abrupt perturbation following gradual learning may appear to be a novel context (even though the rotation size at the end of learning is the same in both conditions), because it has never experienced the large errors all at once in the past. Although this prediction is consistent with at least one prior study^27^, more recent work has exhibited opposing results where gradual learning can improve abrupt learning rates^26,29,30^, possibly by accelerating implicit learning^26^. This raises a key question: why does gradual learning lead to savings in some experiments, but not in others?

Here we consider the possibility that typical sensorimotor learning paradigms may fail to produce savings not due to differences in how individuals learn, but rather because of how the learned behavior decays between the learning and re-learning phases. Specifically, sensorimotor savings is typically studied in paradigms where individuals adapt to a perturbation during an initial learning period, their performance returns to baseline levels during a washout period, and they are exposed to the same perturbation during a re-learning period (in psychological terms, this ‘washout’ period is analogous to ‘extinction’). For visuomotor rotation tasks, conditions employed for washout of initial learning vary across studies. In some cases, a washout period consists of abrupt cessation of the perturbation and includes the same number of trials as the initial learning period^27^. Other studies use a much shorter washout phase^26,29^ consisting of augmented feedback that constrains the cursor motion to an exact straight line (i.e., zero visual error, termed invariant error-clamp^29^) or trials where visual feedback is removed entirely (termed no-feedback trials^26^). This no-feedback condition is akin to moving a hidden cursor on a screen.

Critically, these differences in washout conditions alter the size or quality of errors that induce performance decay. Here we propose that *larger, or more salient,* errors experienced during a washout period might reduce the nervous system’s ability to save what was learned from the initial learning period. In the present study, thus, we examine the overlooked possibility in motor learning, that the nature of washout plays a central role in the expression of savings following gradual learning. Across three groups we manipulate the types of washout errors by (1) removing errors entirely by eliminating visual feedback, (2) limiting the size of errors by removing a visual rotation gradually, or (3) facilitating large errors by abruptly disengaging the rotation. We then apply empirical and model-based approaches to assess whether some washout conditions more easily produce savings during abrupt learning that follows initial gradual learning.

Our current study seeks to not only clarify previous discrepancies in the literature, but also to extend past studies by identifying sources that interfere with the expression of savings. This has important implications for understanding how to preserve increases in our sensitivity to errors^22,26,28^ (thus resulting in greater savings), which in turn may inform new ways to account for motor plasticity in current models of motor adaptation^22,28,31^. Over the long term, this knowledge will facilitate the design of strategies to boost learning rates in neurorehabilitation paradigms.

## Results

Adaptation to a novel visuomotor rotation is often studied in the context of “slicing” movements, in which individuals briskly move their hand (or cursor) *through* a visual target, or point-to-point movements, in which they move their hand to a target and stop on the target. Here, we employed the latter movements that required accurate endpoint control. Our subjects grasped a robotic exoskeleton that controlled the motion of a cursor on a horizontal screen (their arm was hidden from view), and executed point-to-point movements between a start location and one of four targets (Fig. 1a). Initially, the cursor followed the hand’s position in a continuous stroke (baseline), but during the learning periods (exposures 1 and 2), a 24° visual rotation was applied to the cursor’s motion (Fig. 1b). A washout period was placed in between the two learning periods in which the rotation was removed through three different strategies (abrupt removal, gradual removal, or completely removing visual feedback during each movement). Our primary goal was to examine how errors experienced during the washout phase protect or abolish visuomotor savings differently depending on the washout conditions.

**Figure 1.**
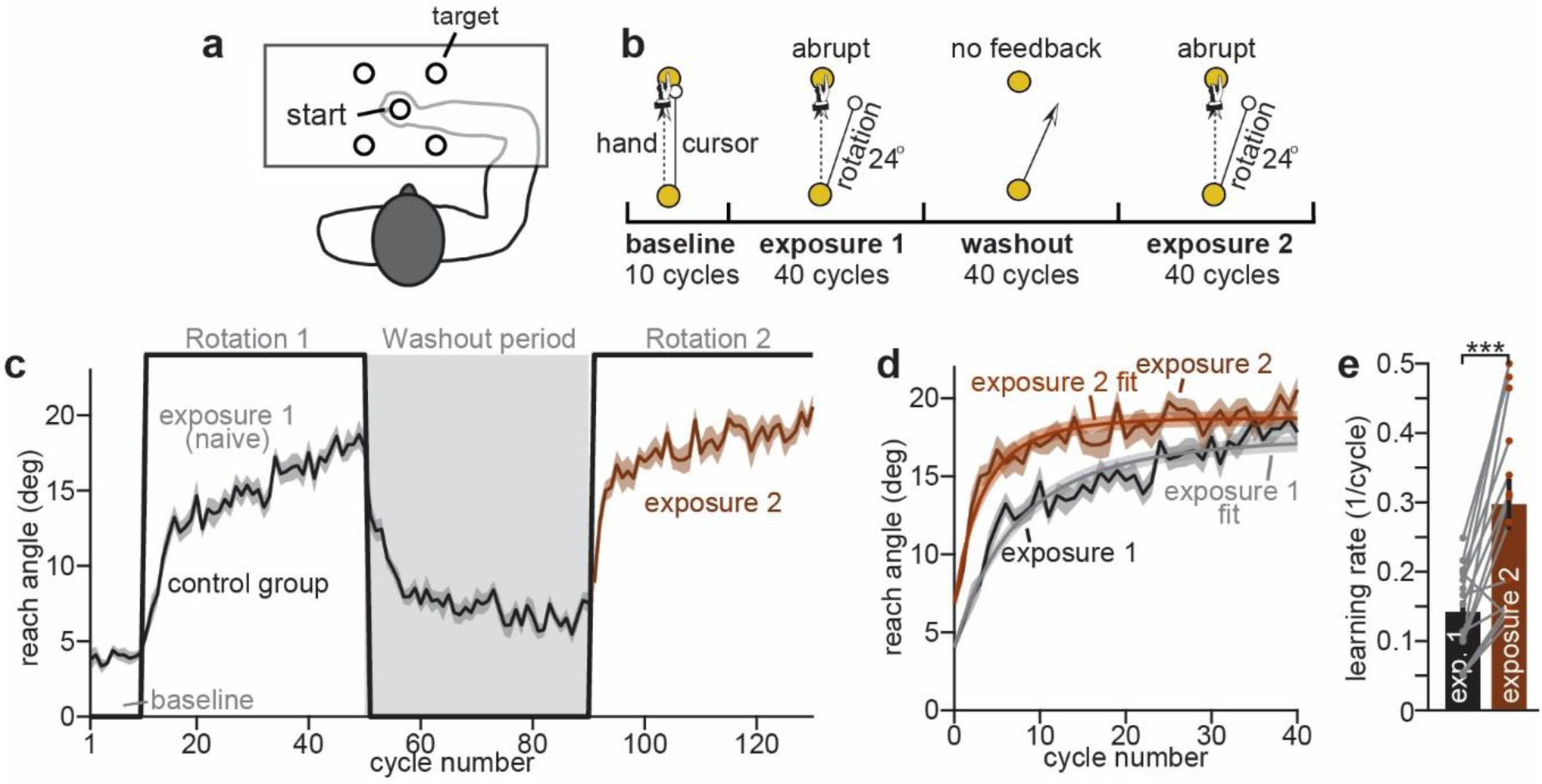
Savings in a point-to-point visuomotor rotation task. **a**. Experiment apparatus. Subjects were seated in a KINARM exoskeleton and executed point-to-point reaching movements to one of four targets. **b**. Perturbation schedule for abrupt adaptation group. This control experiment consisted of 4 experimental periods: (1) 10-cycle baseline period, (2) 40-cycle abrupt 24° rotation (exposure 1), (3) 40-cycle no-visual feedback washout, (4) 40-cycle re-exposure to abrupt 24° rotation (exposure 2). **c**. Learning curves. Solid black line indicates size of rotation during each period. Reach angle at maximum velocity was calculated on each trial and averaged within each 4-trial cycle. **d**. Savings visualization. Learning curves during first and second exposure to the rotation are shown. We fit a 2-parameter exponential curve to individual participants to quantify learning rates. The empirical model fit (empirical fit) is also shown overlaid on the measured learning curves (exposure 1 or exposure 2). **e**. Quantification of savings. We compared the learning rate parameters across exposures 1 and 2 quantified via the exponential model fit in d. Errors bars indicated mean ± SE. Single points in e represent individual participants. Statistics: *** indicates p<0.001.

### Visuomotor adaptation during point-to-point movements is accelerated by visuomotor savings

To confirm that our point-to-point movement design leads to savings (as we recently reported^32^), we first tested a group of subjects (control group) who experienced a visuomotor rotation abruptly during the initial learning period. After a short baseline period, a 24° rotation was abruptly introduced. Over time subjects adapted their reach angles to partially counter the rotation (Fig. 1c, exposure 1). To washout the initial learning, we removed visual feedback, causing reach angles to gradually decay towards their baseline state (Fig. 1c, washout period). To determine whether subsequent learning was improved by prior experience, we abruptly introduced a 24° rotation again, causing a rapid change in reach angle (Fig. 1c, exposure 2). How did learning during the second exposure period compare to naïve adaptation?

To answer this question, we isolated the two learning periods and fit an exponential curve to each participant’s reach angles during both adaptation phases (Fig. 1d). The exponential curve produced an excellent fit to behavior, yielding a root-mean-squared-error (RMSE) within 0.57° of baseline cycle-to-cycle variability (i.e., the theoretical floor for model error: see Methods: *Evaluating model error*; RMSE is shown in Supplementary Fig. 1). As expected, prior exposure to the rotation accelerated adaptation, almost doubling subject learning rates (Fig. 1e; paired t-test, t(14)=5.16, p<0.0001, Cohen’s d=1.33). A power law model for learning, commonly applied for psychological learning^33–35^, also showed an increased learning rate during the second exposure (see Supplementary Fig. 2; Fig. 2a shows model fits; Fig. 2b provides power parameter; paired t-test, t(14)=4.13, p=0.001, Cohen’s d=1.07; model RMSE provided in Supplementary Fig. 1). In sum, we find that point-to-point adaptation to a 24° rotation produces savings and is amenable to further study.

**Figure 2.**
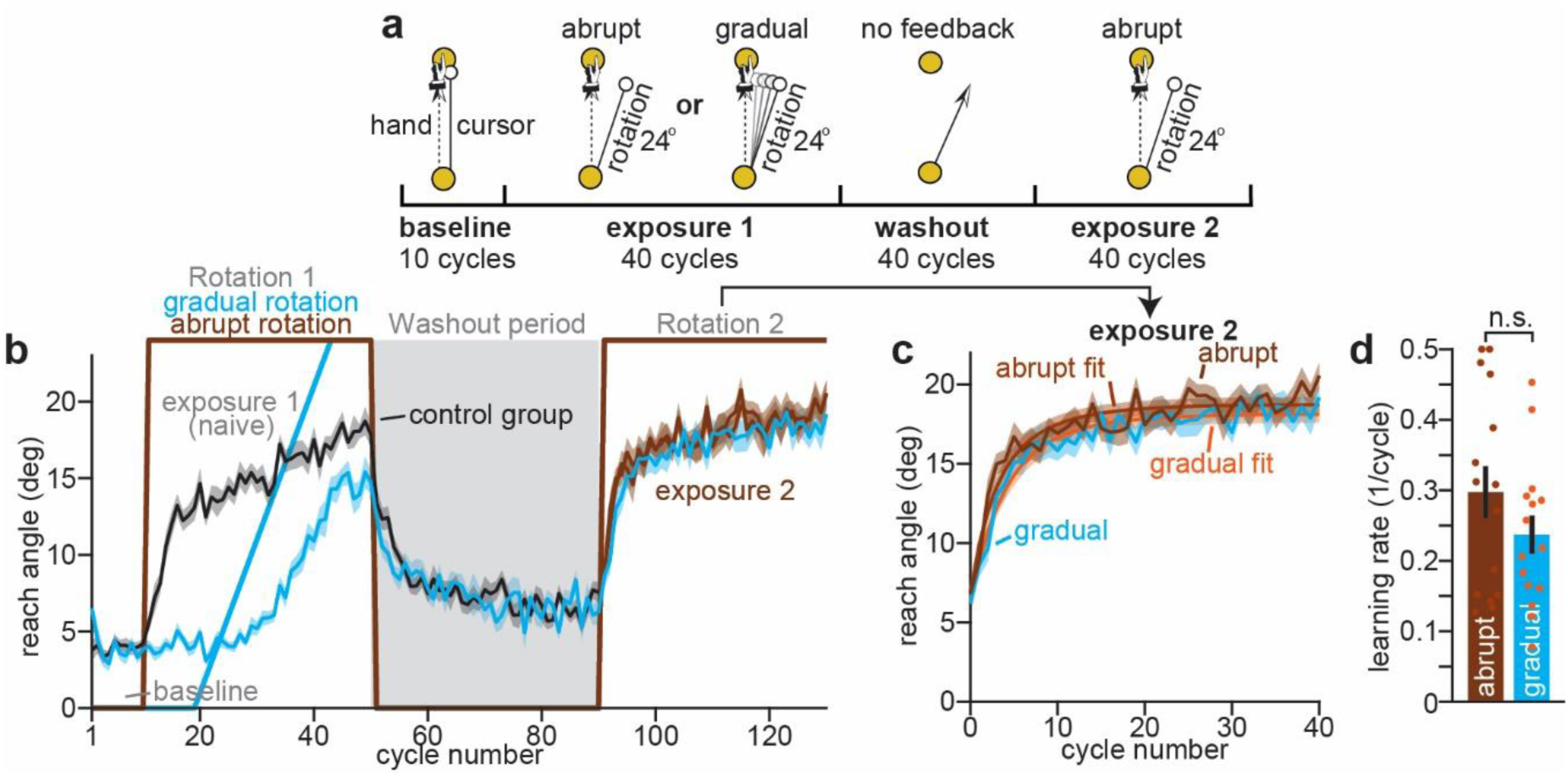
Abrupt and gradual adaptation lead to similar expression of savings in a point-to-point reaching task. **a**. Perturbation schedule for abrupt or gradual adaptation group. Groups differed only based on the visuomotor rotation encountered during the initial exposure: control group from Fig. 1 experienced a 24° rotation that immediately reached its terminal value (labeled abrupt here); gradual group experienced a rotation that slowly increased in magnitude across 24 cycles to reach the same terminal value. **b**. Learning curves. Solid brown line indicates size of rotation during each period. Reach angle at maximum velocity was calculated on each trial and averaged within each 4-trial cycle. Control group (black during exposure 1, and brown during exposure 2) behavior is compared to gradual group (cyan) behavior. Perturbation schedules are overlaid with the data. **c**. Savings visualization. Learning curves during the second exposure period are shown; both groups adapted to an abrupt rotation during this period. We fit a 2-parameter exponential curve to individual participants to quantify learning rates. The empirical model fit (empirical fit) is shown overlaid on the measured learning curves (abrupt or gradual). **d**. Quantification of savings. We compared the learning rate parameters during exposure 2 quantified via the exponential model fit in c. Errors bars indicated mean ± SE. Single points in **e** represent individual participants. Statistics: n.s. indicates p > 0.05. Abbreviations: “gradual” indicates gradual adaptation group; “abrupt” indicates abrupt adaptation group.

### Measuring re-learning rates after initial abrupt and gradual adaptations

Past studies have yielded conflicting evidence as to the necessary conditions for savings. In some cases, savings appears to require the experience of similarly sized errors during both the initial learning and re-learning periods^27^. This is achieved by creating large errors in each learning period via an abrupt perturbation. Other studies, however, have observed savings without initial experience of large errors^26,29^. In this case, the rotation is introduced gradually whereby its magnitude slowly increases during initial exposure to the rotation.

To examine how gradual learning influences subsequent abrupt learning, we tested a second group of participants with a gradual adaptation paradigm (Fig. 2a). Whereas the abrupt adaptation group (controls) in Fig. 1 experienced a 24° rotation all at once during exposure 1, participants in this gradual adaptation group experienced a time-varying rotation that grew 1° per cycle over 24 cycles to reach its terminal value. Linear models fit to the learning curves of the abrupt and gradual adaptation groups (Supplementary Fig. 3) confirmed that gradually introducing the rotation slowed the rate of adaptation (additional participant groups described in the following Results section were used in this analysis; see Methods: *Comparing gradual and abrupt learning rates*; statistics provided in Supplementary Table 1).

After the initial, gradual adaptation (exposure 1), participants in this group experienced a no-feedback washout period, followed by abrupt adaptation (exposure 2). To compare between the gradual and abrupt adaptation groups, we isolated the learning curves during exposure 2 (Fig. 2c) and fit both exponential and power law models to assess their rates of learning. We did not detect a statistically significant difference in re-learning rate between the groups that experienced the rotation gradually or abruptly during initial learning (Fig. 2d shows exponential model; t(28)=1.315, p=0.199, Cohen’s d=0.48; Supplementary Fig. 2d show the power law model; t(28)=0.90, p=0.377, Cohen’s d = 0.33), indicating the presence of savings following gradual adaptation, a finding consistent with prior studies^29^. That being said, this null result may reflect insufficient statistical power. The critical way to confirm whether the initial gradual rotation led to savings is to compare re-learning rates to a naïve control condition. This is pursued in greater detail in the following Results section.

### Size of error during washout modulates the strength of visuomotor savings

It is unclear what causes gradual adaptation to induce savings in some conditions,^26,29,30^ but not others^27^. We considered the idea that savings between initial learning and re-learning phases might be shaped by the washout period that separates the two learning phases.

As shown in Figs. 1 and 2, abrupt and gradual adaptation groups experienced a no-feedback washout condition, preventing the experience of any visual errors during the washout period (Fig. 3b, cyan). To assess whether the washout condition alters the expression of savings, we tested two additional gradual adaptation groups (Fig. 3a) that differed based on the type of error experienced during washout. In the abrupt washout group, the rotation immediately ended at the end of the initial learning period, creating large, oppositely signed errors that induced a rapid decline (Fig. 3b, green). In the gradual washout group, the rotation gradually decreased cycle-to-cycle during the washout period. This prevented the occurrence of large errors in the opposite direction, leading to a gradual decline in performance (Fig. 3b, magenta).

**Figure 3.**
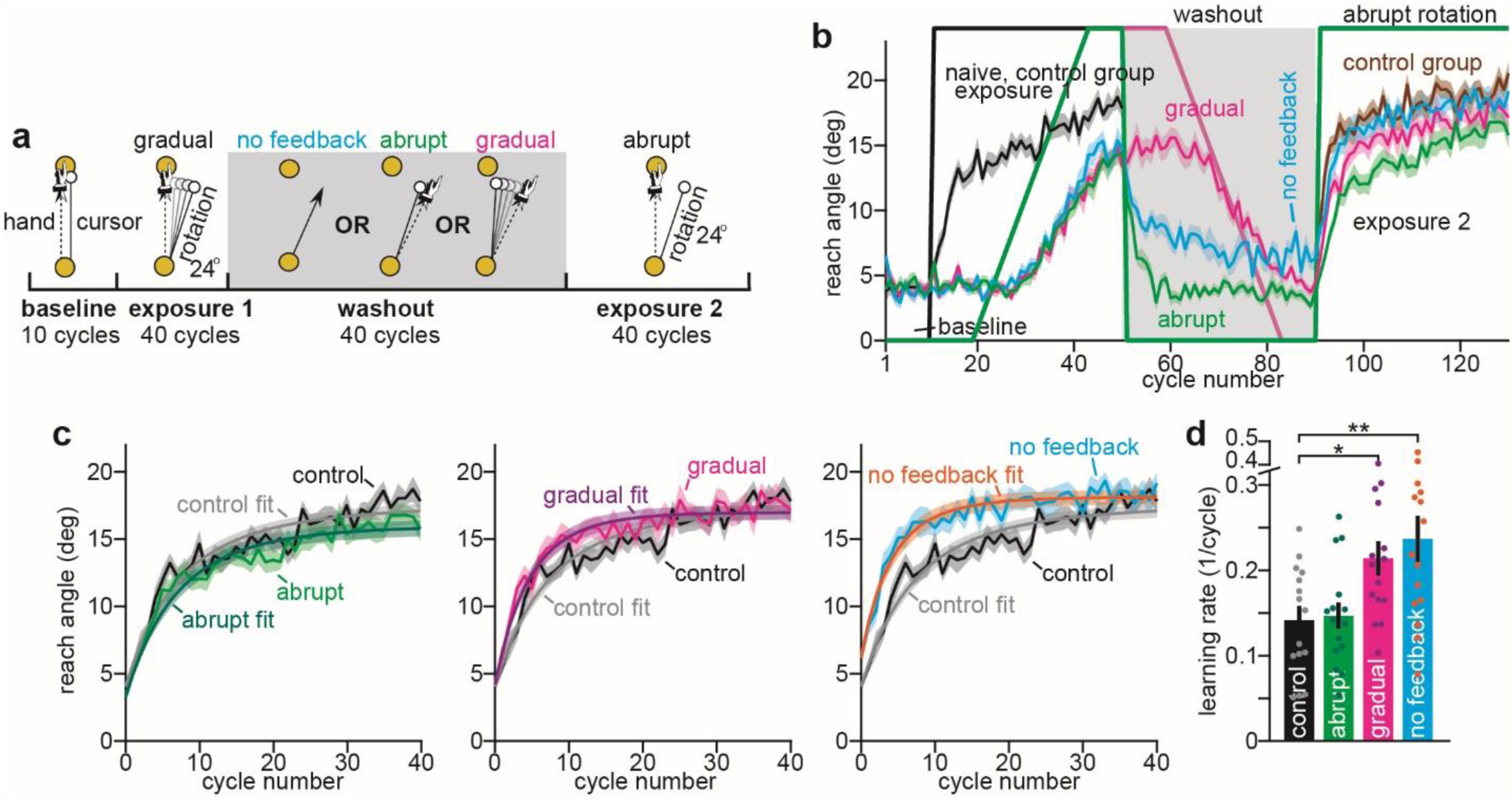
Savings after gradual learning is attenuated by washout error magnitude. **a**. Perturbation schedule for gradual adaptation groups in 3 washout conditions (no feedback, abrupt, gradual). All groups experienced a gradual rotation during exposure 1 and an abrupt rotation during exposure 2. Groups differed only based on errors experienced during washout. One group deadapted in the absence of visual feedback (cyan, no feedback). Second group deadapted in the presence of large errors due to abrupt removal of the rotation (abrupt, green). Third group deadapted in the presence of small errors due to gradual removal of the rotation (gradual, magenta). **b**. Learning curves. Solid lines in different colors indicate size of rotation for corresponding group during each period. Reach angle at maximum velocity was calculated on each trial and averaged within each 4-trial cycle. Abrupt adaptation group (black and brown) behavior is compared to all 3 gradual adaptation groups from described in inset a. The perturbation schedules are overlaid with the data. **c**. Savings visualization. Learning curves during exposure 2 (abrupt adaptation) are shown. Each gradual group is shown in a separate inset (left: abrupt washout; middle: gradual washout; right: no-feedback washout). All learning curves are overlaid with naïve performance of the control group that experienced initial rotation abruptly. We fit a 2-parameter exponential curve to individual participants to quantify learning rates. The empirical model fit (empirical fit) is shown overlaid on the measured learning curves (naïve, abrupt, gradual, or no feedback). **d**. Quantification of savings. We compared the learning rate parameters during exposure 2 in each gradual group to the learning rate parameters in the initial abrupt exposure of the control group. Errors bars indicated mean ± SE. Single points in **d** represent individual participants. Statistics: * indicates p < 0.05.

Following that, all three washout groups were exposed to an abrupt 24° rotation (i.e., exposure 2). To determine how washout influenced subsequent learning rates, we compared reach angles during this abrupt adaptation period to those of naïve participants who had not previously experienced a rotation (*i.e.*, the initial learning period of the control group shown in Fig. 1). Comparing the rates of learning to a naïve control group is a standard approach to detect acceleration in learning rates in conditions where initial adaptation occurred in a gradual fashion^13,26,27,29,36–38^. As before, we fit both exponential and power law curves to participant reach angles to empirically quantify individual subject learning rates (Fig. 3c for exponential model; Supplementary Fig. 2e for power law model). Both the exponential and power models exhibited excellent matches to behavior and possessed an average RMSE within 0.59° of the theoretical minimum error level estimated based on intrinsic cycle-to-cycle reach variability during non-adaptation periods (RMSE distributions are shown in Supplementary Fig. 1; we did not detect a statistically significant difference between the power law and exponential model errors, paired t-test, t(59)=0.71, p=0.478).

Critically, both empirical model types showed that altering washout errors had a dramatic effect on re-learning rates (Fig. 3d, 1-way ANOVA for exponential model, F(3,56)=5.49, p=0.002; Supplementary Fig. 2f for power law, F(3,56)=5.82, p=0.002). Namely, savings was observed only after no-feedback washout (post-hoc Dunnett’s test, p=0.004 for exponential model and p=0.002 for power law) and gradual washout (Dunnett’s test, p=0.04 for exponential model and p=0.045 for power law), but not after abrupt washout (post-hoc Dunnett test, p=0.996 for exponential model and p=0.985 for power law). In sum, initial gradual adaptation did induce savings, and the ability to express savings was modulated by errors subjects experienced during washout. When large errors occurred in our abrupt washout condition, savings was eliminated. When errors were masked (either by making them small with gradual washout, or removing them via no feedback), savings was preserved.

### Improved rates of re-learning are not due to a bias in initial performance state

Improved re-learning in exposure 2, as shown in Fig. 3, could be driven by two different sources: (1) a difference in the starting point for re-learning (i.e., the adapted state each subject possesses at the end of washout; we will refer to this as the ‘terminal washout state’) or (2) an increased rate of re-learning irrespective of the starting point for re-learning. This distinction is critical given that the no-feedback washout group experienced less performance decay during washout (Fig. 4c provides the terminal washout state; 1-way ANOVA, F(3,56)=10.86, p<0.0001) than other experimental groups (post-hoc statistics in Supplementary Table 2), meaning that they started their re-learning period at an advantaged (i.e., larger) learning state.

**Figure 4.**
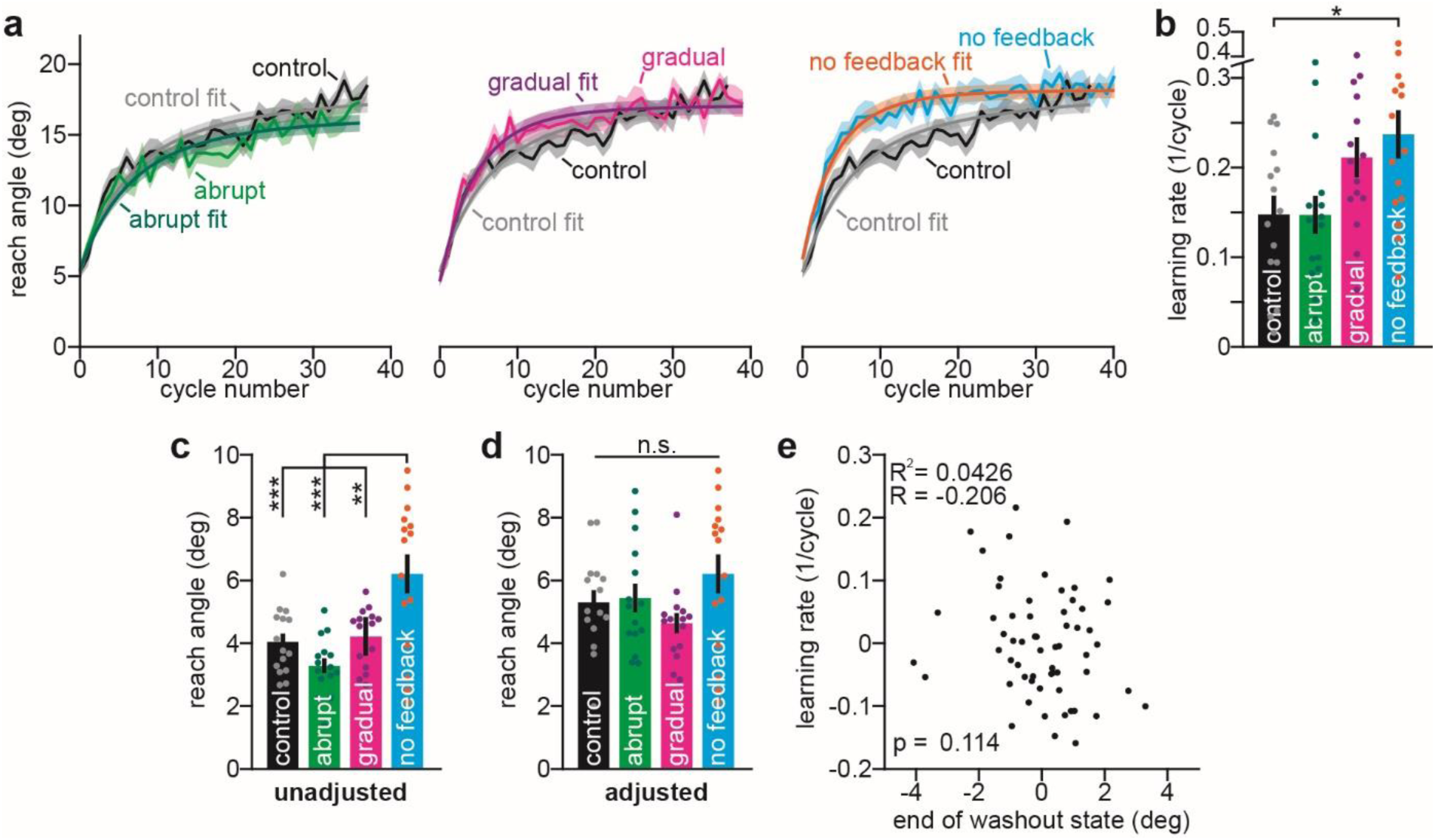
Improved re-learning is not caused by a biased starting point. Exponential and power law models in Fig. 3 were fit to the entire re-learning period in exposure 2. No-feedback washout group started this period at a biased state. Inset **a** (‘unadjusted’) provides the terminal washout state (average angle on last 4 washout cycles (control shows average on last 4 baseline cycles). We refit the exponential model to participant data after removing the initial cycles in the re-learning phase required for Control, Abrupt washout, and Gradual washout groups to ‘catch up’ to no-feedback washout starting point. Inset **d** shows the initial reaching angle on the adjusted starting cycle. The exponential model fits to the adjusted data are shown in Inset **a.** Learning curves in a are overlaid with control group for ease of comparison. The exponential model fit is shown overlaid on the measured learning curve. Inset **b** shows the associated rate parameters for the exponential model. Inset **e**, provides a comparison between re-learning rates (or the naïve learning rate in control group) and the adaptation starting point (i.e., the terminal washout state). X-axis provides the terminal washout state for each participant (the collapsed individual points in **C**). Y-axis provides the learning rates shown in **b**. Both axes were zero-centered at the group level (i.e., the group mean was subtracted from each data point prior to visualization and analysis). Statistics in **e** refer to a linear regression. Errors bars in all insets indicate mean ± SEM. Single dots in **b-e** are individual subjects. Statistics: * indicates p < 0.05, ** indicates p < 0.01, and *** indicates p < 0.001.

Did no-feedback washout produce faster re-learning because other conditions needed to ‘catch up’ to their initial adapted state? To answer this question, we conducted a control analysis where we matched the initial state across groups to that of the no-feedback group (see Methods: *Analyzing relationships between initial state and re-learning rates*). That is, we fit the exponential and power law models to only a portion of the re-learning curve. The starting point for the model fit was chosen to exclude the initial catch-up period where subjects in the control, abrupt, and gradual washout groups fell below the initial state of the no-feedback group. The adjusted curves were defined by the starting points in Fig. 4d; this process removed the initial state bias between groups (1-way ANOVA, F(3,56)=1.94, p=0.134). Importantly, we obtained similar results as before; our model (exponential fit shown in Fig. 4a; power fit shown in Supplementary Fig. 2g) suggested that learning rates during re-learning periods differed between groups (F(3,56)=3.9, p=0.013 for exponential model; F(3,56) = 4.9, p=0.004 for power model), with no-feedback washout exhibiting faster re-learning (post-hoc Dunnett’s test against control, p=0.022 for exponential and p=0.007 for power), a marginal effect in the gradual washout group (p=0.137 for exponential and p=0.079 for power), and no effect in the abrupt washout group (p=1 for exponential and p=0.999 for power).

To corroborate these group-level results, we analyzed the relationship between terminal washout state and re-learning rates at the individual participant level. We collapsed data across all four groups and de-trended datasets (see Methods: *Analyzing relationships between initial state and re-learning rates*) to prevent correlations from being caused by group-level effects. Linearly regressing re-learning rates (Fig. 4e, y-axis) onto the terminal washout states (Fig. 4e, x-axis) did not yield a statistically significant correlation (R=-0.206, p=0.114, R^2^=0.043), and instead trended toward an inverse relationship such that the subjects with greater terminal washout states adapted *more slowly* (counter to the idea that better retention of initial learning speeds re-learning).

In sum, we did not observe clear evidence for the idea that less performance decay in the washout phase would lead to faster re-learning. The improved learning rates we observed in Figs. 3 and 4 appeared to reflect an enhanced capacity for learning as opposed to a bias in the starting point for learning.

### Savings following gradual learning is promoted by an increase in sensitivity to error

Short-term sensorimotor adaptation is due to the combined effects of two processes: (1) trial-to-trial error-based learning, and (2) trial-to-trial performance decay (generally called ‘retention’ or ‘forgetting’ in the sensorimotor learning literature)^15,22,28,39–42^. In principle, savings could be caused by either an increase in the amount of learning that follows the experience of an error, an increase in the amount of learning retained after time passage, or a combination of both sources. These two processes can be captured within a state-space model that represents adaptation as a recursive process controlled by two parameters: an error sensitivity (controls the amount of learning from error) and a retention factor (controls the amount of decay)^15,20,22,28,39,42^.

To determine if the savings we observed following gradual adaptation was due to changes in error-based learning or performance decay, we fit state-space models to individual subject learning curves during the second rotation period in all three gradual adaptation groups (Fig. 5a, green, magenta, blue). To estimate learning and performance decay measures in naïve participants, we fit a state-space model to the initial abrupt adaptation period in the control group shown in Fig. 1 (Fig. 5a, black). To ensure that our state-space model required both a retention parameter and sensitivity to error term, we compared the ‘complete’ model to a simpler one where retention was set to a value of 1 (no performance decay). A model comparison with Akaike’s information criterion (AIC) indicated that the complete model was superior in 55 of 60 participants (see Supplementary Fig. 4a to visualize fits for the reduced model; model comparison is shown in Supplementary Fig. 4b). As such, only the complete parameter model was used in all subsequent analyses. As with power law and exponential model fits, the state-space model yielded a close match to individual subject behavior, with the average error falling within 0.65° of the theoretical minimum value estimated during periods without any adaptation (see Methods: *Evaluating model error* and Supplementary Fig. 1).

**Figure 5.**
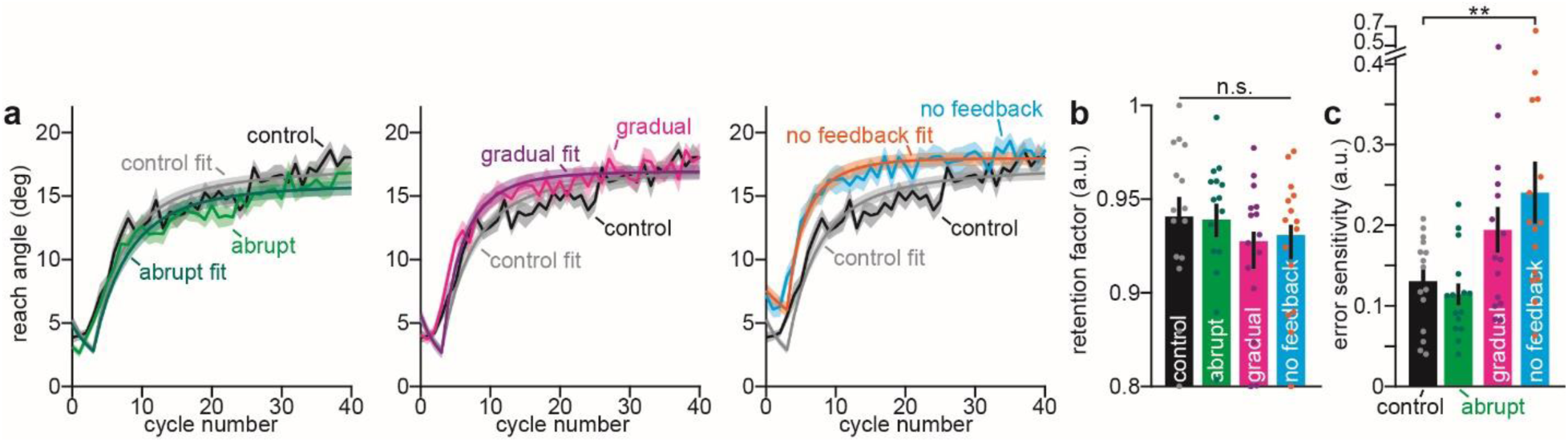
Savings after gradual learning is mediated by an increase in error sensitivity. **a**. Learning curves and visualization of savings. Reach angle at maximum velocity was calculated on each trial and averaged within each 4-trial cycle. Each inset compares naïve adaptation (control, black) to 1 of 3 gradual adaptation groups that experienced different washout conditions (left: abrupt washout; middle: gradual washout; right: no-feedback washout). We fit a state-space learning model (model) to individual participant reach angles that describes learning as a recursive trial-to-trial process of error-based adaptation and decay. **b**. Comparison of retention factors. Retention factors estimated by state-space model fits are compared across the control group during exposure 1 and 3 gradual adaptation groups during exposure 2. **c**. Comparison of error sensitivity. Sensitivity-to-error parameters estimated by state-space model fits are compared across the control group during exposure 1 and 3 gradual adaptation groups during exposure 2. Errors bars indicated mean ± SE. Single points in **b** and **c** represent individual participants. Statistics: n.s. indicates p > 0.05, ** indicates p < 0.01.

To determine if the initial gradual adaptation and subsequent washout period altered the stability of the motor memory during the readaptation period, we compared retention factors across the four groups (Fig. 5b). We did not observe a statistically significant change in retention after exposure to the gradual rotation (one-way ANOVA, F(3,56)=1.43, p=0.244).

To determine if instead, a change in error sensitivity boosted adaptation rates during the second exposure period, we compared the state-space model error sensitivity parameters across the four groups (Fig. 5c). As in past studies^11,16,22^, prior adaptation did alter the sensitivity to error during the second exposure (1-way ANOVA, F(3,56)=6.99, p=0.0004), reflecting the accelerated re-learning rates (Fig. 3d). Interestingly, we found a statistically significant increase in error sensitivity over naïve, in the no-feedback condition (post-hoc Dunnett’s test, p=0.0011), but not in the abrupt condition (post-hoc Dunnett’s test, p=0.994). In the gradual washout condition, we observed a trend towards an increase in error sensitivity over the naïve condition, though this trend did not reach statistical significance (post-hoc Dunnett’s test, p=0.123).

Taken together, this indicates that initial gradual adaptation elevates sensitivity to error, leading to savings. However, errors experienced during the washout phase can reverse this gain in error sensitivity. Enhancement in error sensitivity is optimally preserved when visual errors are totally removed in the washout phase via a no-feedback condition.

## Discussion

The conditions required to increase sensorimotor learning rates are not completely known. This is clearly illustrated in gradual adaptation paradigms where participants adapt to a perturbation slowly and are then re-tested in an abrupt context following a period of washout. Whereas some studies have observed more rapid learning in this situation^26,29^, other accounts have found that gradual learning cannot improve abrupt learning^27^. We considered whether this contradiction is due to the washout phase that separates the two learning phases in savings paradigms. Specifically, does salience of oppositely signed errors during washout inhibit savings after gradual learning?

Here we examined this idea by altering the type of error experienced during the washout phase after gradual learning. We examined no-feedback, gradual, and abrupt washout conditions that differ along an error size gradient. (1) Removing visual feedback eliminates visuomotor error completely, causing the adapted behavior to decay in performance in the absence of overt errors ^43^. (2) With gradual washout, errors are present, but are reduced in magnitude because both the motor action and the perturbation decrease in tandem. (3) Similar to abrupt learning, abrupt washout results in the largest errors due to the immediate removal of the perturbation when subjects still remain at a highly adapted performance level. Our findings suggest that the savings produced by gradual adaptation is protected when washout occurs gradually or in the absence of error, but not abruptly (Fig. 3d).

The idea that washout impacts savings qualitatively aligns with prior work by Kojima et al. (2004),^9^ which demonstrated that savings in the oculomotor system is sensitive to the duration of washout: when the duration of the washout phase is increased, the learning rate during the re-exposure phase reverts to its baseline state, abolishing savings. Thus, both the duration of error presentation and the error magnitude during washout (as shown in our current study) seem to negatively interact with the expression of savings. This hypothesis may explain the mixed results noted above in visuomotor studies of gradual learning. Both Coltman et al. (2021)^29^ and Yin & Wei (2020)^26^ employed short washout periods and observed savings following gradual learning, whereas Herzfeld et al. (2014)^27^ applied longer washout periods and did not find savings after gradual learning. This possibility could easily be studied in future experiments that vary the length of the washout, along with varying types of washout conditions after an initial gradual learning period.

Along these lines, Kitago et al. (2013)^43^ remarked that increasing the number of washout trials appeared to dampen the magnitude of savings relative to other studies that used much fewer washout trials^44,45^. That being said, their observation was made in the context of an abrupt-then-abrupt savings protocol, instead of the gradual-then-abrupt design we used. The savings that follows abrupt or gradual learning may have critical differences. For example, we observed that removing errors in the washout phase after gradual learning protected savings, but the same washout condition abolished savings after abrupt learning in Kitago et al. (2013)^43^ (see Bao et al.^46^ for the case of interlimb transfer).

This discrepancy may be due to variations in the utilization of implicit and explicit learning processes. Gradual perturbations reduce the error size, in turn limiting one’s cognitive awareness of the perturbation. In visuomotor rotation tasks, reducing the awareness impairs cognitive strategies that can be used to counter the rotation, allowing subconscious learning (i.e., implicit) systems to increase their contribution to the adaptation process^22,23,26^. Thus, during the initial *gradual* learning period, we expect greater reliance on implicit adaptation, and reduced engagement of explicit strategies in our tasks, relative to the tasks employed by Kitago et al. (2013). Further, both Kitago et al. and Bao et al. used a single reaching target, whereas our current study incorporated four targets. Increasing the target count enhances task complexity, reducing explicit strategy use and boosting implicit learning (as measured via procedural aftereffect trials^47,48^). Thus, washout may influence savings differently in our study, where the initial learning that is being washed out is largely implicit, as compared to the studies by Kitago et al. and Bao et al., where learning may have relied more heavily on explicit strategies. This idea matches the observation that savings in gradual-then-abrupt learning paradigms like ours have been linked to an enhancement in implicit learning rates^26,30^, whereas savings in abrupt-then-abrupt protocols is due to accelerated explicit responses^7,12^. Thus, we hypothesize that washout has divergent effects on the ability of implicit and explicit systems to save: removing feedback during washout protects savings in the implicit system, but impairs explicit savings.

That being said, it is critical to note that implicit and explicit processes were not measured in our study, and thus, our hypothesis remains both speculative and limited. To examine implicit and explicit contributions to savings in our paradigm, future experiments could assay each system at regular intervals during learning, washout, and re-learning. One approach would be to include implicit probe trials (trials where participants are instructed to abandon any explicit strategy that they are using) to measure the subconscious contribution to their adapted state. Without studies like this, we cannot definitively attribute the observed savings effect to either the implicit or explicit process.

To understand what learning property links washout to the capacity to save, we applied a state-space model that casts short-term adaptation as a cumulative process of trial-to-trial, error-based learning and decay. This model showed that savings relied on an upregulation in error sensitivity, which was reversed by large errors in the washout phase (Fig. 5c). This outcome comports with past work, suggesting that increases in error sensitivity are the primary source of accelerated re-learning after abrupt, gradual, or random perturbations^29^, and the adaptation of disparate motor systems including reaching^16,22,29^ and walking^11^. Intriguingly, our results suggest that increases in error sensitivity are undermined by opposite errors experienced in between learning periods.

We found that error sensitivity was higher in the subjects who washed out without feedback, suggesting that improvements in error sensitivity is protected in the absence of error. We found a trend towards an increase in error sensitivity in the gradual washout group as well (Fig. 4c), but this did not reach statistical significance. Because our exponential model (Fig. 3d) and power law model (Supplementary Fig. 3f) suggested the presence of savings in the gradual washout group, we suspect that our study lacked the sufficient power to detect an enhancement in error sensitivity in the gradual washout condition (power was analyzed using learning rates, see Methods: *Power analysis*). Thus, it remains to be determined whether error sensitivity is reduced by the experience of *any* opposing error, or whether the size of the opposing error amplifies the decline in error sensitivity.

Our findings have critical implications for an influential error sensitivity model defined by a memory of errors^13,27,28^. The memory of errors model that was initially proposed by Herzfeld et al. (2014)^27^ describes changes in error sensitivity based on statistical properties of error. Namely, when two errors have consistent signs, the nervous system could have adapted more to the initial error. A critical model property is locality: consistently small errors or large errors modulate error sensitivity specific to errors of that magnitude. Thus, contrary to our findings, a memory of errors model predicts that a gradual perturbation will not lead to savings during an abrupt perturbation, because abrupt errors are larger than those experienced during a gradual perturbation. Thus, our results add to a growing body of evidence^26,29^ suggesting a need to revisit properties of the error sensitivity landscape; it may be that the experience of consistent errors generalizes more broadly to unexperienced errors of disparate sizes than previously appreciated.

The possibility that changes in error sensitivity extend beyond similarly sized errors could also help to understand declines in learning rate in anterograde interference paradigms^22,42,49–51^. That is, whereas increases in error sensitivity have been linked to improvements in learning when two similar perturbations occur in sequence, a decline in error sensitivity appears to slow learning when transitioning between two opposing perturbations (e.g., clockwise and counterclockwise rotations)^42^. The memory of errors model has not been altered to describe this interference effect. The limited application of the memory of errors model to interference paradigms arises, because learning in response to “positively signed” errors during the first exposure does not produce error sensitivity changes that generalize to “negatively signed” errors during the second exposure. Our findings, that large, oppositely signed errors during washout reverse prior enhancements in error sensitivity (Fig. 5c), suggest that experience of a sensorimotor error produces two distinct effects: (1) an increase in sensitivity to similar errors, and (2) a suppression in error sensitivity to opposite errors. This second suppressive learning property may be a critical process missing from current memory of errors models that would aid in interpreting both the dissolution of savings following certain types of washout, as well as interference effects.

We speculate that antagonistic interaction between the response to positive and negative errors may arise within cerebellar learning circuits that mediate error-based learning, a structure known to encode error directionality based on anatomical connections with the inferior olive^52–54^. That is, an error of a given direction (*i.e.*, sign) leads to complex spiking in certain Purkinje-cell (P-cell) subpopulations in the cerebellar cortex, and suppression in others. Complex spike events in turn lead to long term potentiation or depression at P-cell synapses with parallel fibers^55^. While it is unknown how error sensitivity is encoded by P-cells, our findings suggest the possibility that prolonged absence of complex spikes (when errors are opposite the neuron’s preferred direction) during washout periods may produce synaptic changes that reduce the plastic learning potential of these cells in the future (*i.e.*, during re-exposure). This idea, and the source of cerebellar error sensitivity more generally, remain to be examined.

Note that the state-space model posits that a memory is created during the initial learning period and then decays during washout. Savings reflects the rate at which the original memory can be reacquired. This idea, that savings is related more to a memory of how to respond to errors than to a memory of past actions, is supported by various motor learning studies^13,27,28^. However, the psychology of cognitive decision making as well as subconscious classical conditioning mechanisms, point towards a more nuanced view of learning, namely, as a composition of multiple memories that compete with one another to be expressed^56^. In this view, savings depends on inferring the correct memory to draw upon in the current context.

In general, learning involves predictions. Adaptation is achieved by predicting an outcome and updating one’s actions and beliefs based on errors in this prediction. Broadly speaking, errors encoded in the cerebellum^52,53,57^ are also represented within dopaminergic systems^58,59^ and can be modeled via reinforcement learning algorithms^60,61^. This process is distilled to its fundamental elements in conditioning paradigms, where a stimulus is provided that precedes an appetitive or aversive outcome with some probability. Such experiments have been used extensively to study savings: cases where an initial response is acquired, extinguished, and reacquired within different systems such as eyeblink conditioning, nictitating membrane responses, and salivation in various species including rabbits, mice, and dogs^62–68^. In Pavlovian tasks, faster reacquisition depends on two variables: (1) how strongly the memory created during initial exposure generalizes to the re-exposure period, and (2) how deeply the memory created in extinction (i.e., washout) interferes with the recall process^56^. That is, memories created during the initial conditioning period are not abolished by extinction, but instead remain latent and require the appropriate contexts and cues to promote their re-expression^69,70^.

Many factors determine how well conditioned memories can be recalled, such as similarity between the contexts surrounding the initial acquisition and re-acquisition periods^71,72^, biological significance of the initial learning events^73,74^, and the ability to couple the initial context and the associated memory^75^. Such conditioning rules may help to understand why savings in the abrupt and gradual contexts explored here can be differentially expressed. The large errors experienced during an abrupt perturbation are salient, increasing their biological significance. Further, in tasks with an abrupt-washout-abrupt design, the similarity between the initial learning and re-learning phases is clearly high, promoting savings. This may be why savings in the abrupt-washout-abrupt design can withstand an abrupt washout period with large opposing errors^7,9,22,27^, while less salient gradual errors that are less similar to abrupt errors create memories with increased susceptibility to disruption during an abrupt washout period as we observed in our study (Fig. 3d).

This same framework can be applied to interpret why washing out without error feedback best facilitated savings from the gradual adaptation phase to the subsequent abrupt adaptation phase. That is, washout creates a new pairing between the stimulus (i.e., target presentation in our study) and the appropriate response^76–78^ (e.g., predicting presence of a visual rotation). When the washout phase is experienced in a similar context to the initial exposure, interference occurs. For example, Bouton (1986)^79^ observed that when extinction persists longer than the initial acquisition, savings is completely abolished; with even longer washout durations, reacquisition can further slow and produce a type of retrograde interference^14,50,80,81^. Intriguingly, altering the context of the learner during the extinction phase in ABA paradigms^76,82^ allowed renewal of the initial memory, because the extinction context formed a separate memory that did not exert interference. For our study, the no-feedback condition (in which no cursor or outcome is present on the screen) was the most dissimilar context from the initial learning and re-learning periods (in which full feedback was provided). For this reason, the washout phase creates a memory in a separable context, protecting the initial conditioned response from potential interference.

A related, yet distinct, viewpoint suggests that creating a new context during the washout period is triggered by the salience of prediction errors. When errors are high during washout, this may be a trigger to create a new memory state instead of updating the original state. For example, in Gershman et al.’s study (2013),^83^ subjects in a fear conditioning paradigm more fully extinguished their fear memory, when the extinction (i.e., washout) phase was executed gradually by reducing the probability of the aversive stimulus in a slow continuous manner, as opposed to all at once. The critical idea was that this gradual extinction would encourage the animal to associate the shock with their initial memory, as opposed to an extinction-specific memory. This hypothesis comports with our findings, namely, that when the learned behavior decays gradually or in the absence of feedback, no washout-specific memory state is formed to interfere with the initially obtained memory. This would promote more rapid expression of the initial memory, hence savings.

Multiple memories and statistical inference are used in Bayesian models of sensorimotor learning. These models incorporate both error-based learning and state inference across multiple sensorimotor contexts^84,85^. In single-context memory of errors models, the same adaptive states apply uniformly across different experimental periods, such that any change in learning rate must be due to variation in constitutive learning parameters: sensitivity to error or memory retention. But with multiple contexts, as in the COIN model described by Heald et al. (2021)^84^, the apparent learning rate is influenced by inferring which context is most likely given sensory information. For the COIN model, visuomotor adaptation involves two contexts: a baseline context and a rotation context. Faster learning during the second exposure is caused not by a change in the response to error, but by a greater propensity to expect the rotation given prior training. Our findings have important implications for this inference step, namely, that the errors (*i.e.*, sensory information) experienced during washout alter the inference process: with large opposing errors during the washout period, the nervous system decreases its expectation that the initial rotation state (with its positively-signed errors) will occur again in the future. A broader comparison between memory of errors models and Bayesian frameworks on the effects of washout on future learning remains to be conducted.

## Methods

### Participants

Initially, 10 healthy young adults per group were tested. We then conducted a power analysis to determine the necessary group size to achieve statistical significance with 80% power. The data from these 40 participants (10 per group) were used to estimate effect sizes for this analysis. Our power analysis indicated that a minimum of 13 participants per group were required to achieve 80% power (see *Power Analysis* below for details). Based on this, we collected an additional 5 participants per group, resulting in n=15 per group (60 participants in total). All subjects were right-handed as assessed by the Edinburgh handedness inventory (Oldfield, 1971). The subjects were randomly assigned to the following groups: one abrupt adaptation (i.e., control) group and three gradual adaptation groups. All experimental protocols were approved by the University of Wisconsin-Milwaukee (UWM) Institutional Review Board (IRB). Subjects gave written informed consent prior to participation, which was approved by the UWM IRB in accordance with the Declaration of Helsinki (IRB #16.163).

### Apparatus

Participants were situated in a robotic exoskeleton (KIMARM, BKIN Technologies Ltd, Kingston, ON, Canada) that provided gravitational support to the right arm (all subjects used this arm in the present study). The exoskeleton was positioned so that the arm was hidden underneath a horizontal display (Fig. 1a). To track the hand’s position, a small cursor was projected onto the display, over the subject’s index fingertip. During each trial, the KINARM projected visual stimuli onto the display, so that they appeared in the same plane as the arm. The visual stimuli consisted of a centrally located start circle (2 cm in diameter) and one of four target circles (2 cm in diameter) located 10 cm away from the start target (Fig. 1a). We sampled the hand’s position in the x-y plane at 1000 Hz. Position data were low-pass filtered at 15 Hz, and then differentiated to calculate velocity. Post-processing, analysis, and modeling were conducted in MATLAB R2018a (The MathWorks Inc., Natick, MA).

### Experimental Design

All subjects experienced 4 experiment phases: (1) a baseline period, (2) an initial learning period (exposure 1), (3) a washout period, and (4) a re-learning period (exposure 2) (Fig. 1b). In the baseline period (Fig. 1b, baseline), participants moved their arm to each of the four targets over a 10-cycle period (all four targets were presented once in a cycle). On each trial, continuous visual feedback was provided via a cursor indicating the location of the index fingertip. Participants were instructed to move rapidly and accurately to the target location. The trial ended 1.5 sec after target presentation. To begin the next trial, the participants brought their hand back to the central start position. Throughout the entire experiment, the four targets were presented in a pseudorandom order in 4-trial cycles.

During the initial learning period (exposure 1), a 24° rotation was introduced abruptly for the abrupt adaptation (control) group (see Fig. 1), and gradually for three gradual adaptation groups (see Figs. 2-5). More specifically, the visual display of the cursor in the control group was rotated 24° counterclockwise (CCW) about the start circle from trial 1 and remained constant throughout the entire exposure period. In the gradual adaptation groups, the rotation size started at 0° for the first 9 cycles, then increased by 1° (CCW about the start circle) per cycle for next 23 cycles, and remained at 24° for last 8 cycles. During the re-learning period (exposure 2), all participants experienced the same abrupt 24° rotation condition as that experienced by the control group in the initial learning period. All participants performed reaching movements over 40 cycles (i.e., 160 trials) in each of the two exposure periods.

In the washout period, the control group performed reaching movements without visual feedback. The three gradual adaptation groups experienced one of three washout conditions: (1) abrupt washout, in which the rotation was completely removed from trial 1 (i.e., the same as the baseline period); (2) gradual washout, in which the rotation size was decreased gradually from 24° down to 0°); and (3) no-feedback washout, in which subjects performed reaching movements without visual feedback. All participants performed reaching movements over a 40-cycle period. In sum, all groups re-learned in an abrupt context. Thus, the four groups were separated by their initial learning and washout conditions. The abrupt adaptation group (controls) is the group that initially learned abruptly and washed out without feedback. The gradual adaptation/gradual washout group is the group that initially learned gradually and then washed out gradually. The gradual adaptation/abrupt washout group is the group that initially learned gradually and washed out abruptly. The gradual adaptation/no-feedback washout group is the group that initially learned gradually and washed out without feedback.

### Behavioral data processing

Data analysis was conducted in MATLAB (MathWorks, Natick, MA). To quantify performance, we calculated the reaching movement angle at peak velocity (c.f., direction error^37^). This angle was computed as the hand’s angular position relative to a line segment connecting the start and target positions. Outlying reach angles were detected and removed as detailed in *Outlier Detection* prior to additional analysis. Next, data were averaged within each 4-trial cycle.

### Comparing gradual and abrupt learning rates

To compare learning rates between gradual and abrupt adaptations, we analyzed learning curves in the exposure 1 period with a linear model. For gradual learning, the linear model incorporated a delay parameter that allowed the starting point at which learning begins to vary between each participant (the parameter was allowed to vary between 9 and 20 cycles: note that 9 cycles were chosen because the rotation was held at 0° for 9 cycles). The model was fit to individual subject behavior and the delay parameter that minimized squared error was selected. Because abrupt learning is not well-captured by a linear model (see Fig. 1c) only the initial 8 cycles were used to quantify an ‘early’ learning rate to compare to the gradual condition. No delay was allowed in the linear fit. Model fits are shown in Supplementary Fig. 3a, and learning rates (i.e., linear slope) are provided in Supplementary Fig. 3b. Abrupt exposure to the rotation more than doubled learning rates (1-way ANOVA, F(3,56)=17.54, p < 1e^-7^; all post-hoc comparisons between abrupt group and each of the 3 gradual learning groups via Tukey’s test yielded p<0.0001; other comparisons p > 0.97). See Supplementary Table 1 for post-hoc t-test statistics.

### Exponential and power law models

To empirically compare adaptation between groups, we assessed the learning rate during each abrupt rotation period via an exponential function. We focused on rate measures, to prevent our evaluation from being biased by the initial state of the learner (*i.e.*, subjects may demonstrate greater total adaptation not because they learn more rapidly, but because their initial reach angle is greater during the second abrupt rotation due to incomplete washout from the initial rotation). Thus, we fitted an exponential curve to participant hand angles, and used the rate parameter as our measure of learning rate. For this, we employed a 2-parameter exponential function designed to enforce that each curve begin at the initial pre-rotation reach angle. For the initial rotation in the control group, this hand angle was computed as the average angle over the last 4 baseline cycles. For the abrupt rotations during the second exposure period (across all groups), this angle was computed as the average angle over the last 4 washout cycles. The exponential function took the mathematical form:

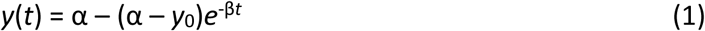

Here, α is parameter that encodes the exponential curve’s asymptote, *y*_0_ is the initial reach angle described above, β is the rate parameter of interest, *y*(*t*) is the reach angle on cycle *t* (starting at *t* = 0). The function was fit to 41 cycles of data at a time (40 cycles of the abrupt rotation plus the initial reach angle cycle on *t* = 0) using a bounded least-squares approach (MATLAB *fmincon*). The parameter search was repeated 100 times, each time varying the initial parameter estimate that seeds the algorithm. The α parameter was bounded between -45° and 45°. The β parameter was bounded between 0 cycle^-1^ and 0.5 cycle^-1^. The model parameters that minimized squared error across all 100 repetitions were selected.

Exponential models are commonly used in sensorimotor learning studies to assess adaptation rates. For learning in the psychological domain, models with a positive hazard such as a power law are more common^33–35^. Thus, to corroborate our exponential model, we repeated our primary analyses using a 2-parameter power law given by:

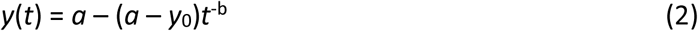

Similar to our exponential model, the power function was constrained to begin at the terminal washout state (*y*_0_) by starting with *t* = 1. In this way *a* sets asymptotic learning and *b* is the power parameter that we used to quantify and compare learning rates. As with the exponential model, *fmincon* was used to fit the model to individual participant data in a space where *a* was bounded between 0 and 45° and *b* was bounded between 0 and 5.

### Analyzing relationships between initial state and re-learning rates

We conducted a set of control analyses (results shown in Fig. 4) to determine how the initial state at the start of the re-exposure period altered learning rate. Note that this initial state at the start of exposure 2 is synonymous with the terminal washout state (as the washout directly precedes exposure 2). Two analyses were conducted: one at the group-level and one at the subject-level.

For our group-level analyses we repeated our exponential model fit to a smaller segment of the re-exposure period (or the initial exposure period in the control group). We selected this period based on the no-feedback group. This group exhibited greater lingering adaptation at the end of the washout phase (see Fig. 4c), meaning that they would begin adaptation at the start of exposure 2 at a higher level than the other groups. To control for this, we identified new starting points for exponential model fits that more closely matched the mean starting point exhibited in the no-feedback group (6.203°). To do this, we used a 2-step approach. First, we used exponential model fits to the entire exposure period (as described in *Exponential and power law models*) and identified when these models crossed the threshold value of 6.203° for each participant. We then trimmed the data to remove all data points prior to this threshold value, such that the beginning of the adjusted learning curves corresponded to the cycle just prior to the threshold crossing (the resulting initial states are shown in Fig. 4d). We then refit the exponential model to the truncated data, using the exact same process as described in *Exponential and power law models*. Resulting learning rate parameters are reported in Fig. 4b). This process was repeated (using the same start points) for the power law model, and results are shown in Supplementary Figs. 2g and 2h.

Finally, we compared learning rates during the re-exposure period (or naïve exposure for the control group) to their terminal reach angle at the end of washout (or the end of the baseline period for the naïve control group). This was done to test the hypothesis that the subjects who had an initial bias towards the adapted state might exhibit an unfair advantage in their learning rate. For this analysis, the exponential rate parameters were compared to the average reach angle at the end of washout (the 4-cycle period described in *Exponential and power law models*). For this, data were collapsed across the four groups (n=60 participants in total). Data were de-trended at the group level by zero-centering; the group mean was subtracted within each group prior to analysis. The resulting data (Fig. 4e) were then compared across participants via linear regression.

### State-space learning model

To interpret our empirical results using an error-based learning framework, we employed a state-space model^15,22,28,39,40,42^. The model describes adaptation as a trial-by-trial learning process whereby the adapted state of the individual is updated by (1) learning from the error experienced on each trial and (2) a small amount of decay that reflects temporal instability of short-term adaptation^41,86,87^. In the model, the total update in the motor memory due to error is controlled by one’s sensitivity to error (*b*) and the strength of memory retention between trials is encoded by a retention factor (*a*). Together, learning and performance decay (this is termed ‘forgetting’ or ‘retention’ in the sensorimotor adaptation literature) collectively determine how the subject’s internal state (*x*) evolves in response to error (*e*) experienced on trial *t* in the presence of Gaussian state noise (ε_x_, normally distributed with zero mean, and standard deviation of σ_x_) according to:

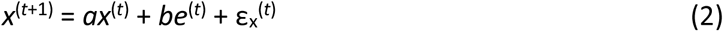

Eq. (2) allows us to ascribe performance differences during the adaptation period to interpretable quantities: retention (*a*) and error sensitivity (*b*).

Note, however, that the internal state (*x*) is not observable. Instead, on each cycle, the motor output (reach angle) is measured. The reach angle (*y*) directly reflects the subject’s internal state, but is altered by execution noise (ε_y_, normal with zero mean, std. dev. = σ_y_) according to:

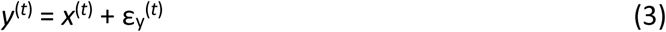

Together, Eqs. (2) and (3) represent a single module state-space model. We fit this model to each participant’s reach angles during the adaptation period using the Expectation-Maximization (EM) algorithm^39^. For each adaptation period, all 40 abrupt rotation cycles and 3 preceding baseline or washout cycles were included. EM is an algorithm that conducts maximum likelihood estimation in an iterative process. We used EM to identify the parameters {*a*, *b*, *x_1_*, σ_x_, σ_y_, σ*_1_*} that maximized the likelihood of observing the data (*x_1_* and σ *_1_* represent the subject’s initial state and variance, respectively). We conducted 100 iterations, each time modifying the initial parameter guess that seeded the algorithm. The retention factor (*a*) was bounded between 0.85 and 1. The initial state (*x*_1_) was bounded between -10 and 20°. Error sensitivity (*b*) was bounded between 0 and 1. Last, all 3 variance parameters were bounded between 1.0e^-3^ and 100. We selected the parameter set that maximized the likelihood function across all 100 iterations.

Last, we conducted a control analysis to assess if a reduced model without performance decay (i.e., complete memory retention; *a* = 1) was sufficient to explain behavior. The same fitting procedure described above was used to fit the state-space model, but the retention parameter was constrained to a value of 1, thereby reducing model complexity by one parameter. The resulting model fits are provided in Supplementary Fig. 4a. Akaike’s Information Criterion (AIC) was used to compare the complete model to the reduced model in Supplementary Fig. 4b.

### Outlier detection

As noted above in *Behavioral data processing*, outlier reach angles were removed prior to model fits, empirical analyses, and statistical testing. To remove outliers, we used a 2-step process which started by eliminating trials where the absolute reach angle exceeded 60° (0.567% of trials). Then, in the second pass, we removed additional outlier trials in each cycle. For this, we collapsed reach angles across all subjects and 4 trials in each cycle, and removed trials that deviated by more than 5 median absolute deviations from the cycle median (1.798% of trials). This was done separately within each experimental group, such that each cycle removal process used 60 trials (4 trials in a cycle, multiplied by 15 participants in the group).

In a control analysis, we employed an alternative way to detect outliers within individual participants without considering other subjects in the group. For this, we binned reach angles in consecutive 10-trial bins. In each bin we calculated the median reach angle. Outliers were taken as trials that deviated from the median by more than a threshold level. We varied this threshold to explore more liberal and conservate outlier detection; 4 levels were used (from most to least aggressive: 15°, 20°, 25°, and 30°) leading to removal of 0.68% to 2.6% of trials. Resulting analysis was done based on our exponential model learning rates (Supplementary Fig. 5). We found that all thresholds produced the same qualitative result (statistics are given in Supplementary Tables 3 and 4), which also matched the conclusions reported in our Results section.

### Evaluating model error

The goodness-of-fit for each model (exponential, power, and state-space model) was quantified by calculating the root-mean-squared-error (RMSE) for each individual participant learning curve during the abrupt learning period (exposure 2 was used for the initial gradual learning groups but exposure 1 was used for the control group). All RMSE measures are shown in Supplementary Fig. 1. For comparison, we also calculated the intrinsic variability in cycle-to-cycle reach angle (shown as a horizontal line in Supplementary Fig. 1). This intrinsic variability provides a theoretical lower bound for the RMSE that could be achieved by any model, as it represents the inherent random movement noise that corrupts movement angles during baseline periods without any adaptation. To calculate this value, we calculated the standard deviation in angle during the baseline period as well as the last 10 cycles of the washout period, and averaged these two values together (note that in the no-feedback group, we used only the baseline standard deviation because during the washout period the rotation had not been removed long enough to achieve a stable steady-state at which to assess motor noise in the absence of learning).

### Statistical analysis

For statistical comparisons between 2 groups or conditions, paired or independent two-sample t-tests were employed. When more than 2 groups were compared, we applied one-way ANOVAs. In post-hoc tests on savings-related measures (see *Measuring savings* below), we used Dunnett’s test to measure differences between each experimental group and the control. For cases where initial learning rates and terminal washout states were compared across groups, Tukey’s test was used to evaluate all possible pair-wise comparisons.

### Measuring savings

Our principal analyses concern whether experiencing a perturbation gradually during initial learning produces a benefit for re-learning in the future. For sensorimotor adaptation, savings refers to an increase in the rate or extent of re-learning. However, in many cases, including our current study, initial learning rates during gradual learning cannot be observed due to a mismatch between perturbation types in exposure 1 and 2. In exposure 1, participants adapt to a gradual rotation. In exposure 2, the rotation is abrupt. The gradual and abrupt learning rates are not comparable: by design, gradual learning is necessarily slower than abrupt learning due to the presence of smaller errors^28,88,89^ throughout the learning process which are known to drive smaller trial-to-trial updates in reach angle^28,88,89^. In our case, abrupt learning was more than twice as rapid as gradual learning (see *Comparing gradual and abrupt learning rates* and Supplementary Fig. 3).

In cases such as ours where initial learning is gradual and re-learning is abrupt, savings is assessed by comparing learning rates during the abrupt re-learning period to that of a separate control group that only experiences an abrupt rotation (this control condition allows us to obtain a proxy estimate for the naïve abrupt learning rate)^13,26,27,29,36–38^. This is the approach we utilized herein, and it informed our choice of statistical tests described in the previous section.

### Power analysis

To determine our group size, we conducted a power analysis. As noted before, we recruited 10 participants in each group and used these preliminary results in our analysis (n=40 subjects total). For our analysis, we selected our principal investigation in Fig. 3. Exponential models were fit for exposure 2 in three experimental groups (groups that experienced gradual adaptation during exposure 1) and exposure 1 in the naïve control condition. The fitting procedure is described above in *Exponential and power law models*. We then extracted the learning rate for each participant and calculated the mean and variance for each group. For each group, we drew simulated learning rates from a normal distribution, and then conducted a 1-way ANOVA. Power was measured based on the finding that p < 0.05 for a given simulated experiment. The resulting power is reported in Supplementary Fig. 6 as a function of group size. Note that 13 subjects were required in our simulations to reach 80% power.

## Data availability

The datasets used and/or analyzed during the current study will be made available upon request by contacting the corresponding author.

## Code availability

The MATLAB scripts used for empirical analyses, model fitting, model comparison, and statistical testing will be made available upon request by contacting the corresponding author.

## Author contributions

MJ, BM and JW designed the experiments and methodology. MJ and BM collected data. MJ and STA analyzed data and created figures. MJ and STA wrote manuscript. MJ, STA, and JW edited the manuscript.

## Acknowledgements

This work was supported in part by the National Institute of Child Health and Human Development (T32HD040127 to STA), the W.M. Keck Foundation (to STA) and a Burroughs Wellcome Fund Careers at the Scientific Interface Award (to STA).

## Supplementary Figures

**Supplementary Figure 1.**
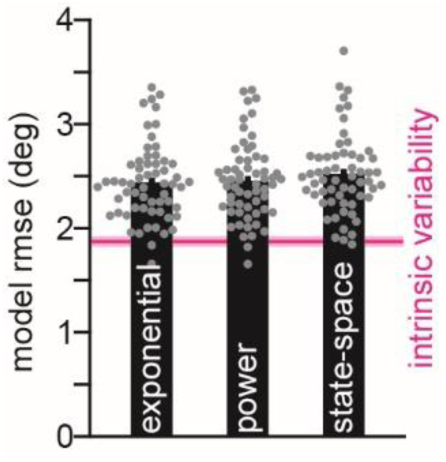

**Supplementary Figure 2.**
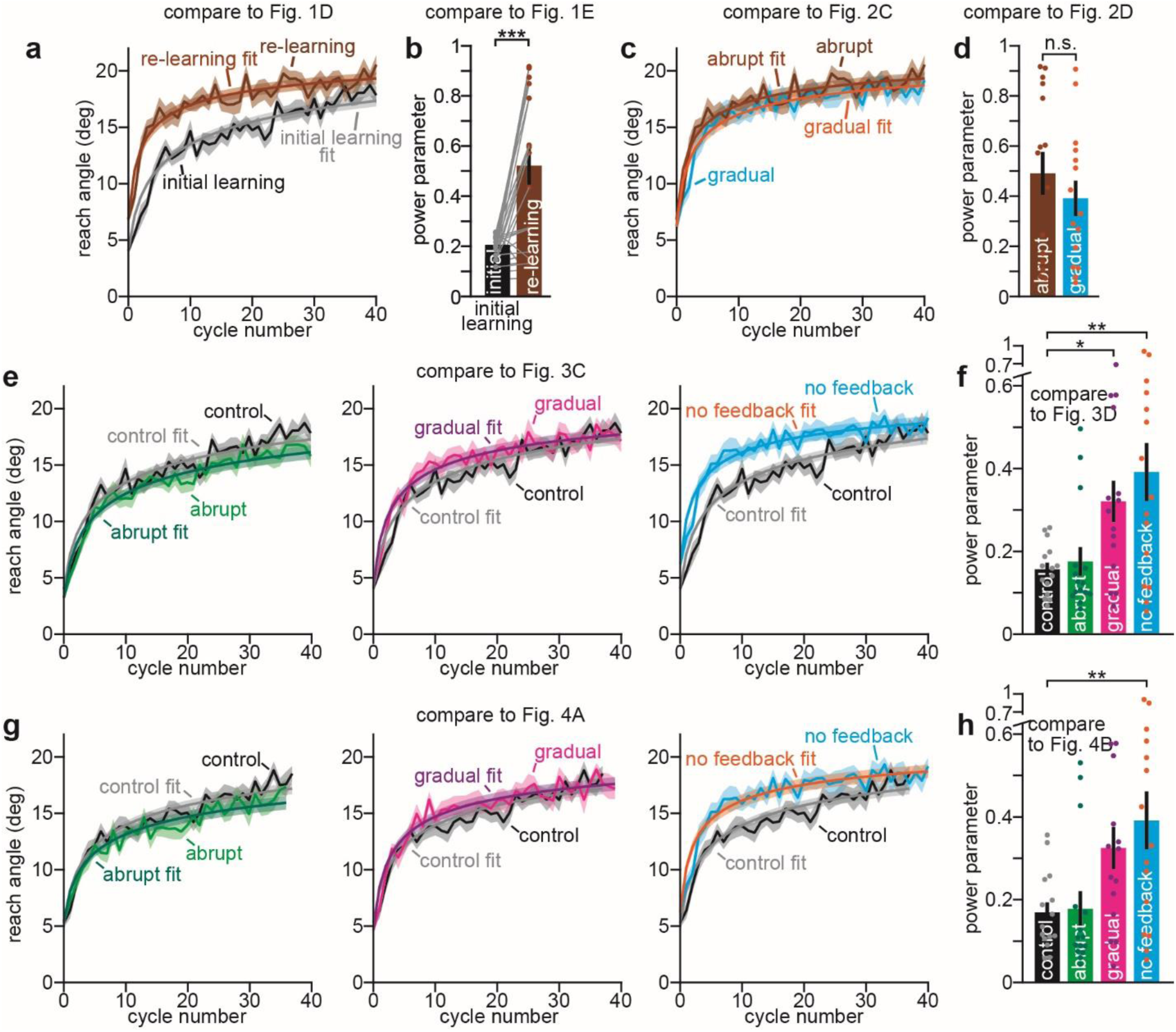

**Supplementary Figure 3.**
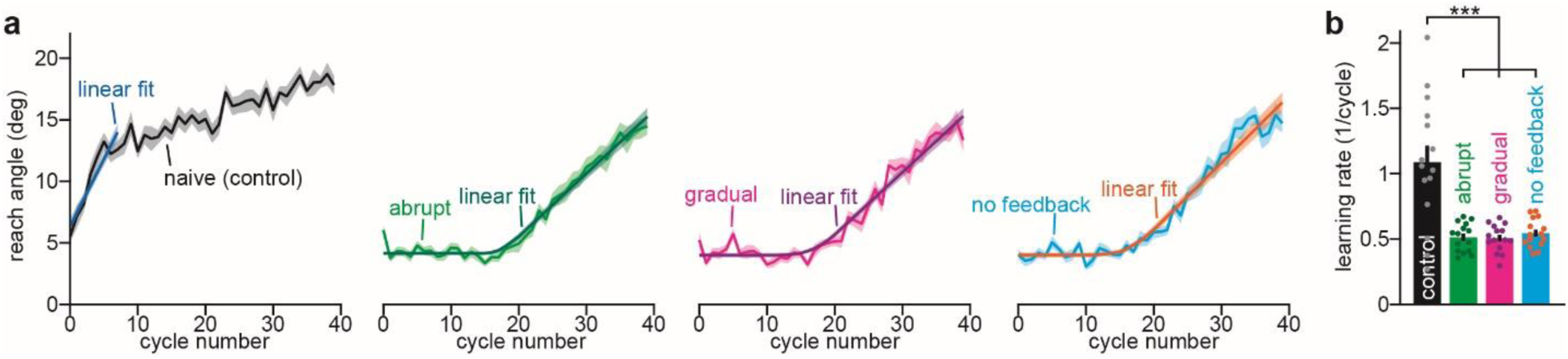

**Supplementary Figure 4.**
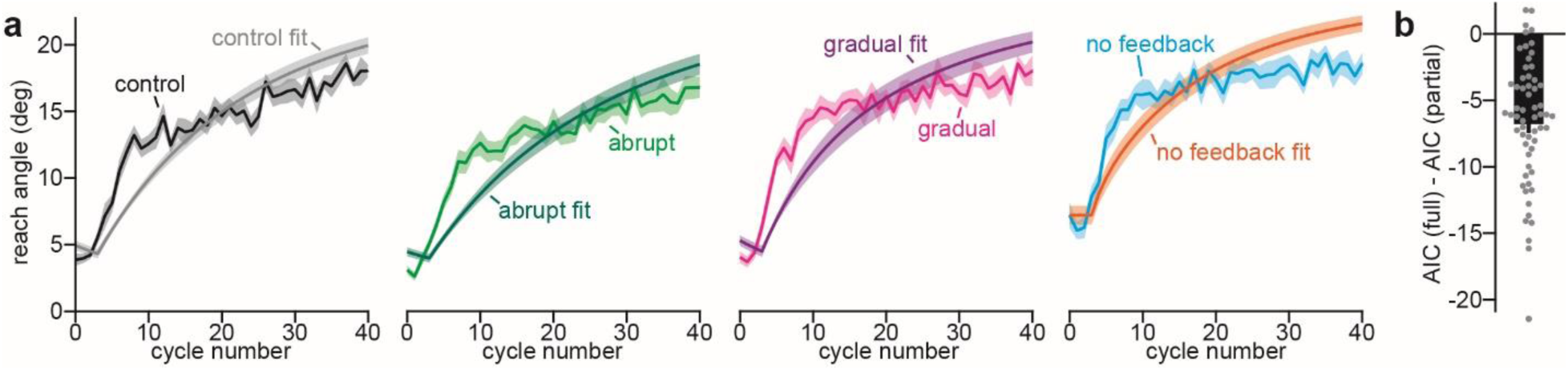

**Supplementary Figure 5.**
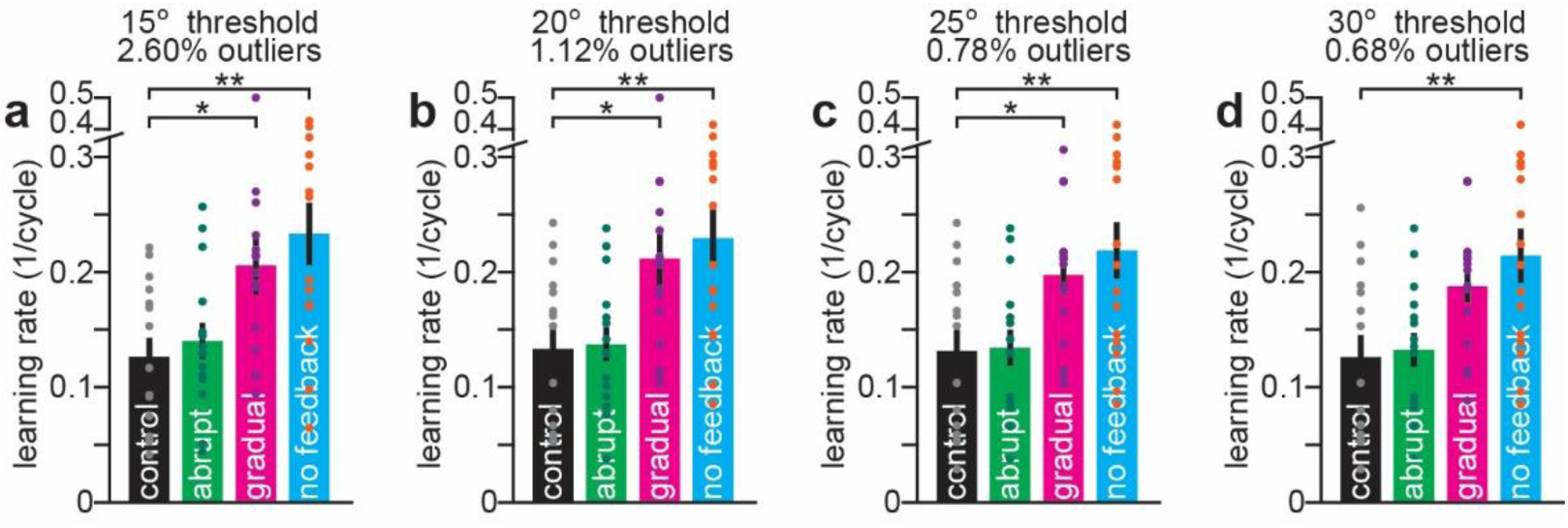

**Supplementary Figure 6.**
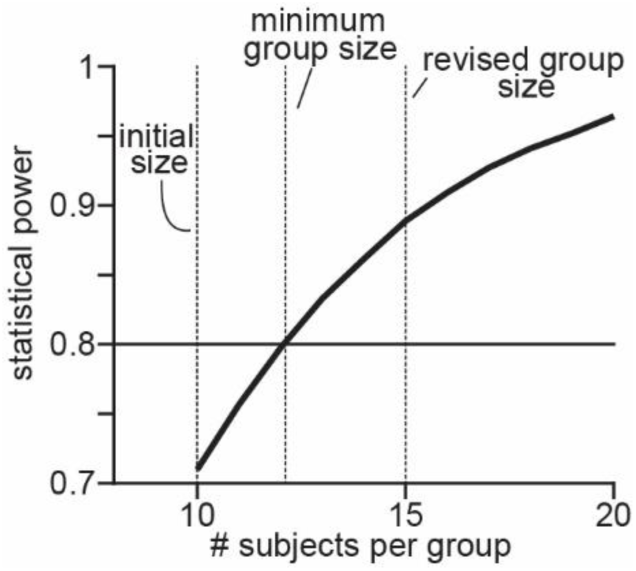

